# Pharmacological arginine deprivation renders pancreatic cancer cells susceptible to senolysis and inhibition of the integrated stress response

**DOI:** 10.64898/2026.08.03.740778

**Authors:** Laura Mainz, Mohamed A. F. E. Sarhan, Lisa Schlicker, Stefanie Kirner, Jennifer Morton, Elena Gerhard-Hartmann, Andreas Rosenwald, Ursula Eilers, Christina Schülein-Völk, Werner Schmitz, Apoorva Baluapuri, Martin Eilers, Elmar Wolf, Markus Diefenbacher, Almut Schulze, Mathias T. Rosenfeldt

## Abstract

Pancreatic ductal adenocarcinoma (PDAC) is almost inevitably fatal and largely resistant to current chemotherapy regimens. Senolysis, i. e. the selective elimination of senescent cells, and the targeting of cancer specific cellular metabolism are two novel treatment approaches that have proven to be effective for a variety of cancer types in pre-clinical studies. We now demonstrate that pharmacological depletion of arginine with pegylated recombinant human arginase 1 (PEG-rhARG1) induced senescence and activated the integrated stress response (IRS) in PDAC cells. Cells treated in such a manner were strikingly susceptible towards senolysis with ABT-263 (navitoclax) and also sensitive towards inhibition of the IRS. These results demonstrate a novel mechanism for an induced sensitivity of PDAC cells that could be exploited for cancer therapy.

## Introduction

The most frequent form of pancreatic cancer, pancreatic ductal adenocarcinoma (PDAC), is essentially not amenable to current treatment [1]. Increasing evidence suggests that the exploitation of specific metabolic requirements of PDAC cells could be a promising treatment concept [1]. Arginine is a semi-essential amino acid, i. e. it can be produced endogenously, but must be supplemented from extracellular sources during high demand [2, 3]. Many cancer cells, including PDAC cells, have an impaired intrinsic ability to generate arginine, due to defects in the enzymatic machinery of the urea cycle, including argininosuccinate synthase 1 (ASS1) and ornithine transcarbamylase (OTC) (Figure 1A) [2, 4]. This reliance on extracellular arginine, termed auxotrophy, imposes a potential vulnerability towards arginine deprivation [3]. Two arginine-depleting agents are currently in clinical trials and are well tolerated [2, 3, 5]. ADI-PEG20 (pegylated arginine deiminase) converts arginine to citrulline and ammonia, but its bacterial origin can elicit an immune response [2, 5]. In contrast, PEG-rhARG1 (pegylated recombinant human arginase 1, BCT-100) is not immunogenic and converts arginine to ornithine and urea [2, 5]. It has been shown that arginine withdrawal causes a G1- or S-phase cell cycle arrest in different cancer cell lines and is associated with enhanced activity of senescence-associated β- galactosidse (SA-β-Gal) in glioblastoma multiforme cells [5, 6]. Senescent cells are characterized by a tumor-suppressive durable proliferation arrest, but also secrete mediators of chronic inflammation, a phenomenon referred to as senescence-associated secretory phenotype (SASP) and that is believed to be the major driver of the pro-tumorigenic effect of senescence [7]. Due to their ambivalent nature it is highly desirable to selectively eliminate senescent cells in the context of cancer therapy. This can be brought about by a group of mechanistically diverse compounds, called senolytics, some of which are already in clinical trials [8, 9]. The integrated stress response (ISR) is another adaptive cellular process that promotes cell survival but can, depending on circumstances, also induce cell death. Targeting the ISR is currently under investigation for numerous cancers [10].

**Figure 1:**
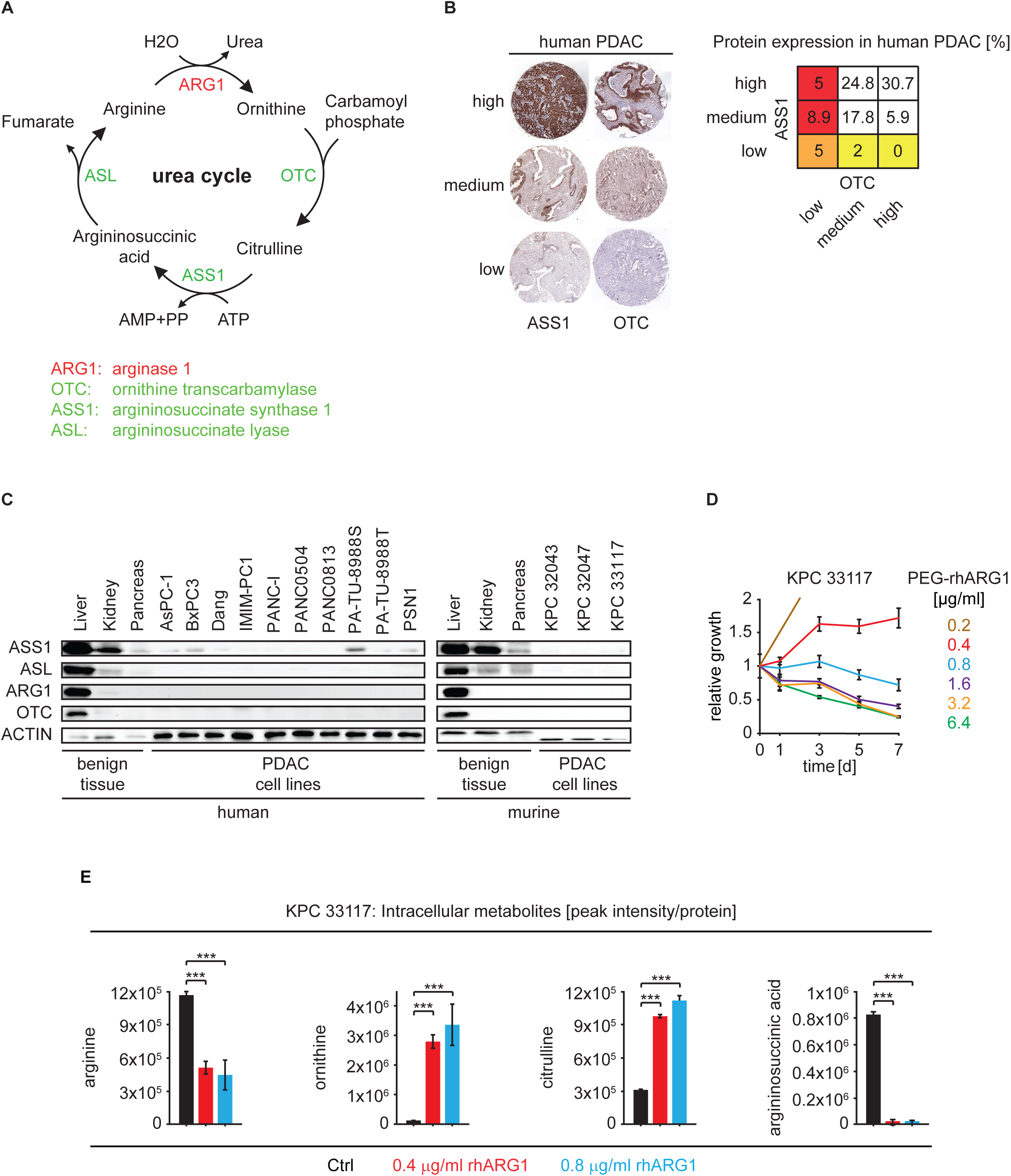
PEG-rhARG1 treatment reduces cell growth. **A)** Simplified urea cycle. **B)** Protein expression of ASS1 and OTC in a cohort of human PDAC patients with exemplary immunohistochemistry. **C)** Immunoblotting of ASS1, ASL, ARG1 and OTC in different tissues and cell lines. **D)** Relative cell growth upon different doses of PEG-rhARG1. **E)** Relative intracellular abundance of selected metabolites of the urea cycle after PEG- rhARG1 treatment for 3d. Statistics: Student’s t test. **P < 0.01, ***P < 0.001.

Here we demonstrate that pharmacological arginine depletion via PEG-rhARG1 renders PDAC cells strikingly susceptible towards senolysis with the BH3-mimetic ABT-263 (navitoclax) and also towards inhibition of the IRS. These results establish a novel mechanism for an induced metabolic sensitivity of PDAC cells that could be exploited to treat a disease for which there is currently no effective therapy.

## Results

### PEG-rhARG1 impairs cell growth and alters the metabolic profile of PDAC cells

ASS1 and OTC are required for the production arginine (Figure 1A). Immunohistochemistry on a tissue microarray (TMA) with 101 individual PDAC specimen revealed low expression of either ASS1 (7%) or OTC (18.9%) in a substantial percentage of patients, implying arginine auxotrophy in approx. 25% of PDAC cases (Figure 1B). Additionally, enzymes of the urea cycle (ASS1, ASL, ARG1 and OTC) were expressed at only minimal levels in human PDAC cell lines compared to benign tissue (Figure 1C). Murine PDAC cells, derived from a well-established mouse model (KPC, *Pdx1-Cre; KRas^G12D/WT^; Trp53^R172H/WT^*) that closely recapitulates human PDAC [11, 12], also lacked expression of these enzymes (Figure 1C).

Treatment with PEG-rhARG1 induced sustained cell growth arrest over a 7-day time course in murine PDAC cells (Figure 1D). Higher doses (above 0.8 µg/ml) also resulted in a reduction of cell number, indicating cell death. For further experiments, we focused on the growth inhibitory effects of PEG-rhARG1 and used the two lowest concentrations that robustly impaired proliferation (0.4 and 0.8 µg/ml). Intracellular metabolite levels were assessed using liquid chromatography-coupled mass spectrometry (LC-MS) in murine PDAC cells treated with either 0.4 or 0.8 µg/ml of PEG-rhARG1 for 3 days. 75 metabolites were significantly altered between control cells and PEG-rhARG1 treated cells, with the higher dose inducing a more pronounced response (Supplementary Figure 1). Intracellular arginine levels were reduced to about 45% of controls. Levels of citrulline and especially ornithine were strikingly elevated following PEG-rhARG1 treatment, while argininosuccinic acid was essentially absent (Figure 1E). Further experiments focused on understanding the nature of PEG-rhARG1 induced growth arrest.

### Arginine withdrawal induces senescence in pancreatic cancer cells

PEG-rhARG1 treated KPC cells consistently exhibited reduced cell growth and BrdU-incorporation (Figures 2A, 2B). Proliferation in the presence of PEG-rhARG was restored in argininosuccinic acid supplemented medium (0.4 mM) (Supplementary Figure 2A). Due to the prolonged nature of the arrest, we hypothesized that arginine depletion might induce senescence. Growth retarded cells featured enhanced activity of SA-β−Gal, chromatin remodeling as evidenced by the occurrence of high mobility group box 3 (HMGB3/HMG2A) foci and tri- methylation at the lysine 9 residue of the histone H3 protein (H3K9me3), elevation of cyclin dependent kinase inhibitor 2a/p16 (CDKN2A/P16) protein, and increased mRNA expression of components of the SASP (Figures 2C-E). Similar results were obtained in human PDAC cell lines (IMIM-PC1 and PA-TU-8988T) (Supplementary Figures 2B-D). Having established a senescent growth arrest upon PEG-rhARG1 treatment, we wondered how the transcriptional profile is affected in arginine depleted cells.

**Figure 2:**
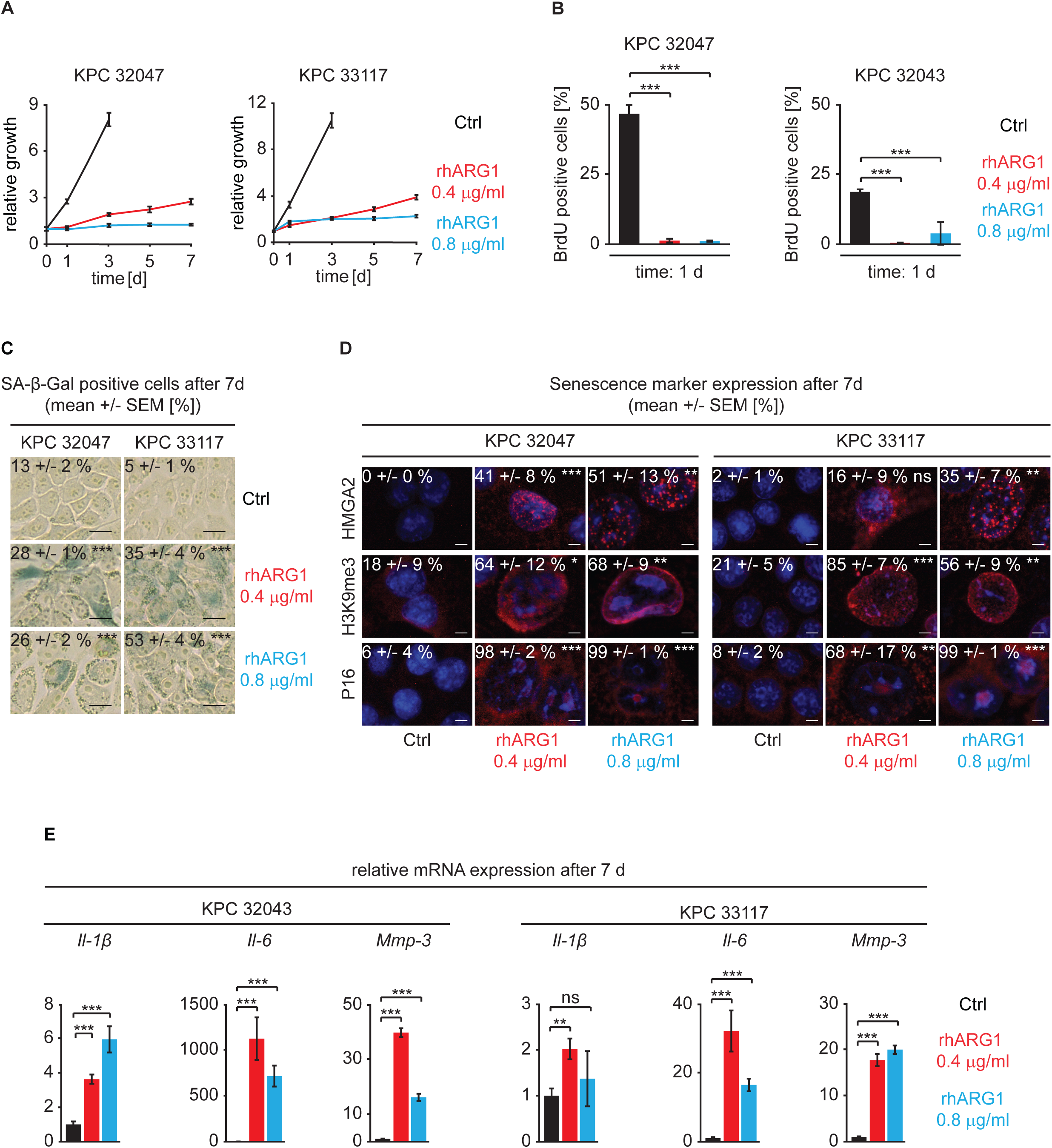
Arginine withdrawal induces senescence. **A)** Relative cell growth upon PEG-rhARG1 treatment. **B)** Quantification of BrdU-incorporation upon PEG-rhARG1 treatment. **C)** Sa-β-Gal activity as indicated. **D)** Immunofluorescence as indicated with quantification. **E)** Relative mRNA expression levels of SASP components. Statistics: Student’s t test. *P < 0.05, **P < 0.01, ***P < 0.001.

### PEG-rhARG1 treatment activates the ISR

RNAseq analysis coupled with gene set enrichment analysis (GSEA) revealed upregulation of the hallmark unfolded protein response (UPR) gene set upon treatment of KPC cells with 0.8 µg/ml PEG-rhARG1 (Figure 3A, 3B). Arginine depletion was accompanied by phosphorylation of the eukaryotic translation initiation factor 2α (eIF2α) and enhanced protein expression of C/EBP-homologous protein (CHOP) (Figure 3C). Likewise, human PDAC cells showed evidence of ISR activation upon arginine depletion (Supplementary Figure 3). We next investigated whether arginine depletion exposed exploitable vulnerabilities.

**Figure 3:**
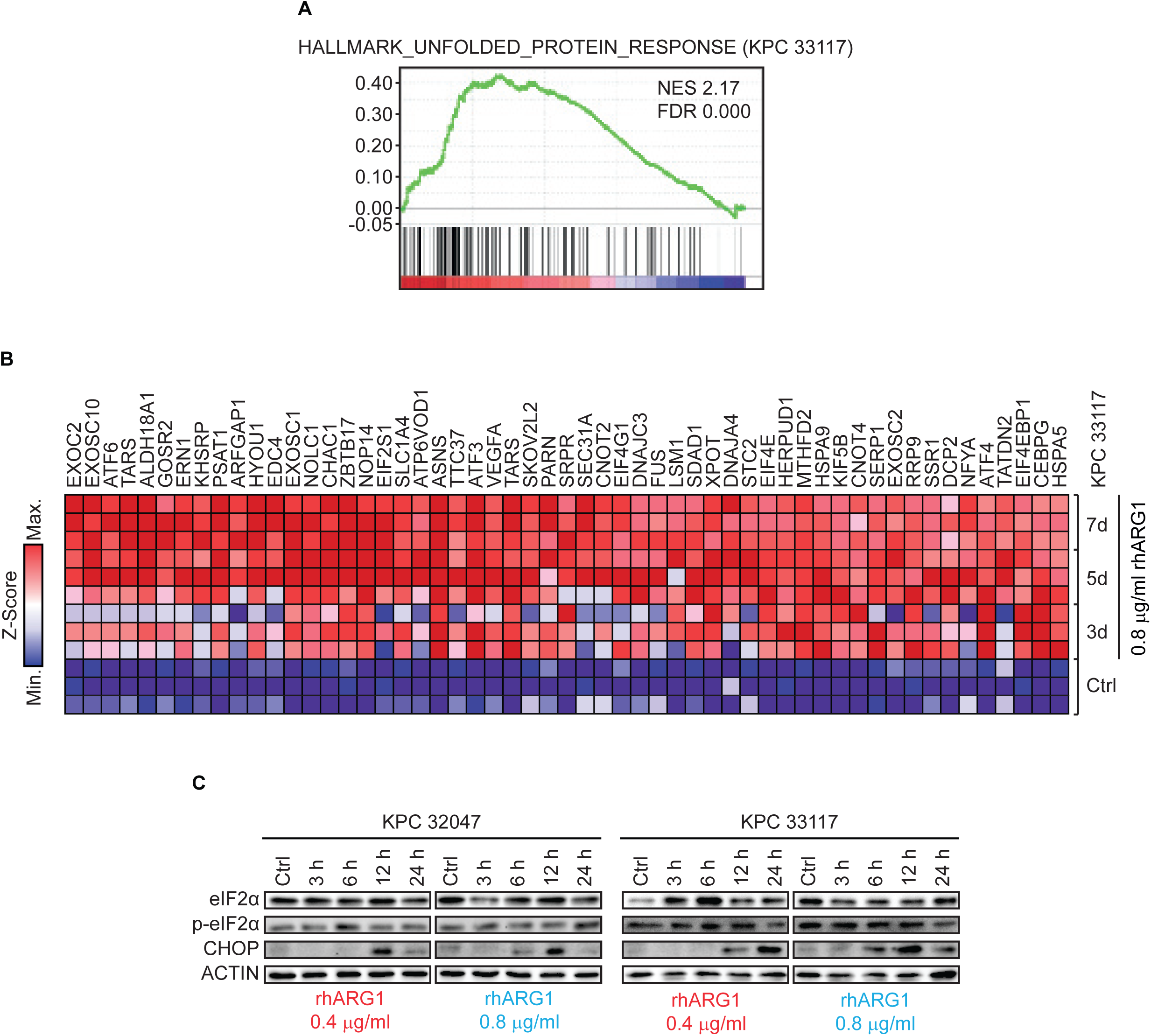
PEG-rhARG1 treatment activates the ISR. **A)** Gene set enrichment analysis of Hallmarks of Unfolded Protein Response in KPC cells upon treatment with rhARG1 for 7 d. **B)** Heatmap of time-resolved Unfolded Protein Response gene expression as indicated. Data presented as Z-score values. **C)** Immunoblotting of eIF2α, p-e2Fα, CHOP and ACTIN as indicated.

### PEG-rhARG1 treated PDAC cells are susceptible to senolysis or inhibition of the ISR

While the senolytic compound ABT-263 alone did not affect proliferation, a combination with PEG-rhARG1 resulted in profound cell death (Figures 4A-C). This was at least partially due to the induction of apoptosis, as evidenced by the cleavage of caspase 3 (Figure 4D). Eukaryotic translation initiation factor 2 alpha kinase 4 (EIF2AK4, GCN2) mediates activation of the ISR upon amino acid starvation [10]. A92, a specific inhibitor of EIF2AK4, only had a minor effect on cell growth when used alone. However, when combined with PEG-rhARG1, a markedly reduced cell growth at least in part due to apoptosis was observed (Figures 4A-C, E). Similar results were obtained in human PDAC cells. Notably, monotherapy with A92 already had a profound antiproliferative effect in PA-TU-8988 cells, but the synergism with PEG-rhARG1 was retained (Supplementary Figure 4). These data reveal a metabolic vulnerability of PDAC cells upon arginine depletion that sensitizes these cells towards senolysis or inactivation of the ISR.

**Figure 4:**
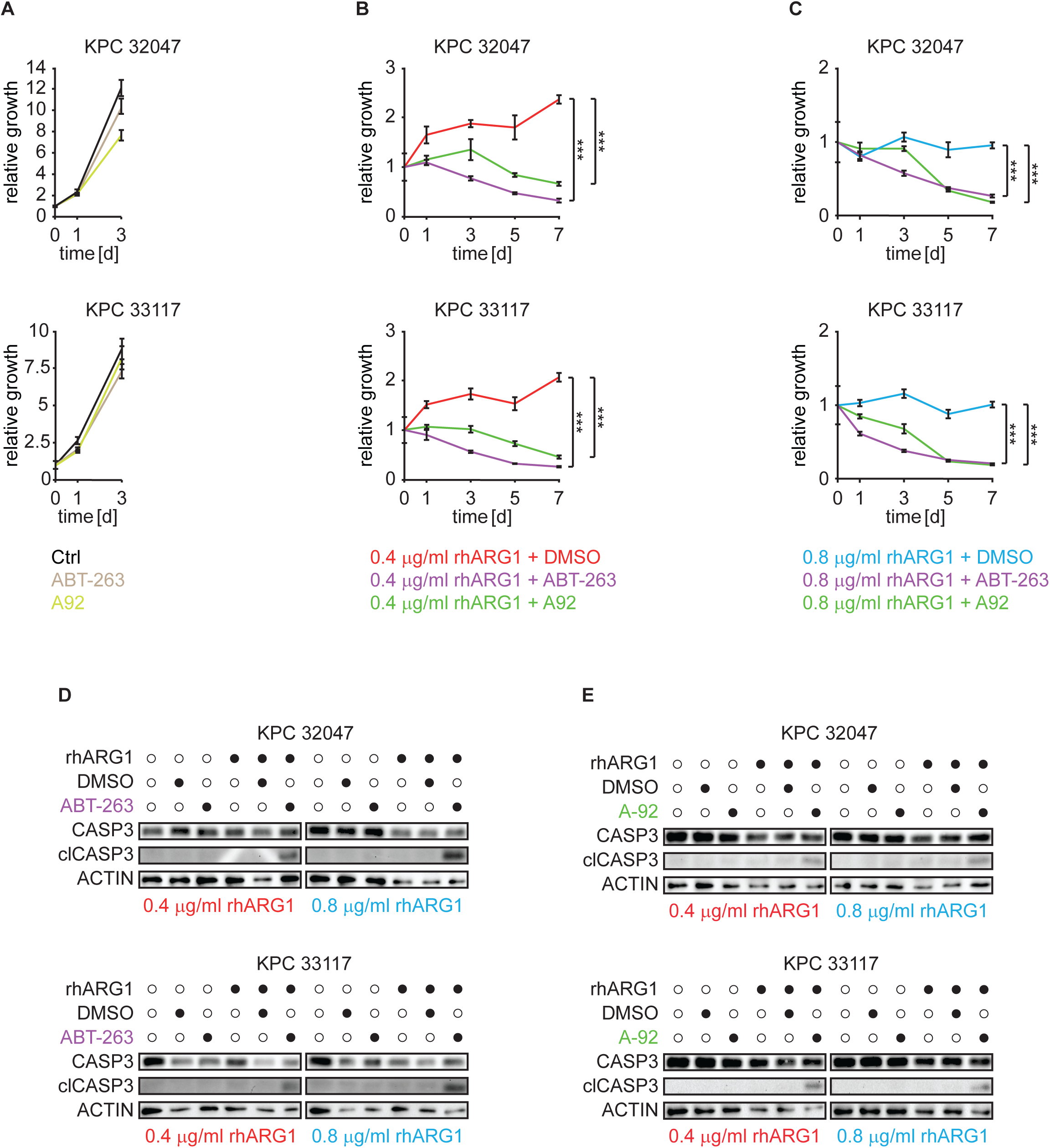
PEG-rhARG1 treated PDAC cells are susceptible to senolysis and to inhibition of the ISR. A-C) Relative cell growth in response to the indicated treatments. **D & E)** Immunoblotting for CASP3, clCASP3 and ACTIN as indicated. Statistics: Student’s t test. ***P < 0.001.

## Discussion

Here we have shown that pharmacological depletion of arginine with PEG-rhARG1 induced senescence and activated the ISR in PDAC cells. Arginine-deprived cells were susceptible to senolysis with ABT-263 and to inhibition of the EIF2AK4/GCN2-mediated ISR. Cells from several cancer types rely on exogenous arginine provision owing to reduced activity of ASS1 or OTC that are required for endogenous arginine production [3]. In agreement with our findings, reduced expression of ASS1 in pancreatic cancer has been reported before and it was shown that PDAC cell proliferation is reduced in response to PEG-ADI treatment, but induction of senescence or the ISR were not demonstrated [13–15]. PEG-ADI arrested glioblastoma cells display enhanced activity of SA-ß-Gal, but other markers of senescence were not investigated [6]. In different cellular systems, arginine depletion caused DNA damage, nucleotide depletion, activation of the UPR and alterations in NF-κB signaling, all of which have the potential to induce senescence [7, 9, 16, 17]. Thus, it is likely that the same mechanisms also contribute to senescence execution in our system.

Although arginine-depleting monotherapy has shown some clinical efficacy, there is increasing evidence that combinatorial therapy holds more promise [2, 5]. It has been reported that certain chemotherapeutics, radiotherapy, or inhibitors of histone deacetylase enhance the effects of PEG-ADI [4, 15, 17, 18]. A recent report found that inhibition of GCN2 in arginine-deprived hepatocellular carcinoma promotes a senescent phenotype that imparts a vulnerability to senolysis, provided GCN2 is impaired as well [16]. Notably, in our system inhibition of GCN2 was not a pre-requisite for susceptibility towards senolysis (or vice versa). While the exact causes for the differences between hepatocellular carcinoma and PDAC remain unclear, the different modes of arginine depletion in both studies could be responsible. We employed a system for the rapid pharmacological conversion of arginine into ornithine to reduce arginine levels. In contrast, in the study by Missiaen et al. arginine levels were reduced via dietary means or by impaired cellular uptake that did not result in concomitant elevated ornithine levels [16]. Strict dietary control of arginine intake might be difficult to achieve in human cancer patients, making pharmacological intervention a more feasible modality that warrants further clinical evaluation.

Our study indicates that pharmacological arginine depletion in combination with senolysis or inhibition of the ISR might hold therapeutic promise in PDAC patients. Importantly, the route to clinical application could be short, as both arginine depleting enzymes and senolytics have undergone clinical evaluation with acceptable systemic side effects [2, 8].

## Materials and Methods

Materials and Methods are provided as supplementary information.

## Supplementary Figure Legends

**Supplementary Figure 1:**
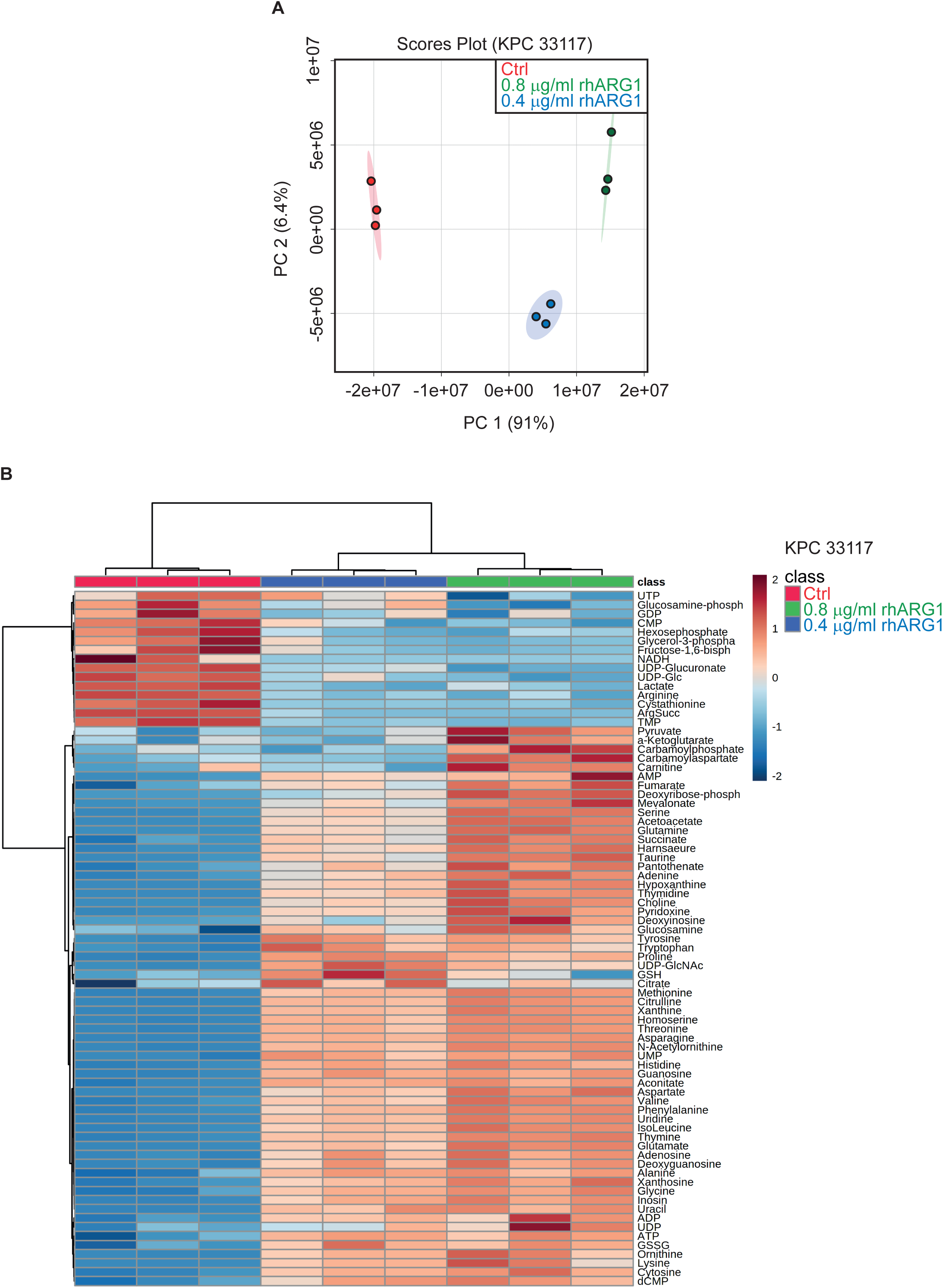
Altered intracellular metabolites after PEG-rhARG1 treatment. **A)** Scores plot comparing polar metabolites extracted from murine PDAC cells treated with 0.4 or 0.8 µg/ml PEG-rhARG1 and controls. **B)** Heatmap showing significantly changed metabolites in murine PDAC cells following treatment with either 0.4 or 0.8 µg/ml PEG-rhARG1 for 3 d. Significance was determined using 2-way ANOVA (p≤0.05). Hierarchical clustering was performed using MetaboAnalyst.

**Supplementary Figure 2:**
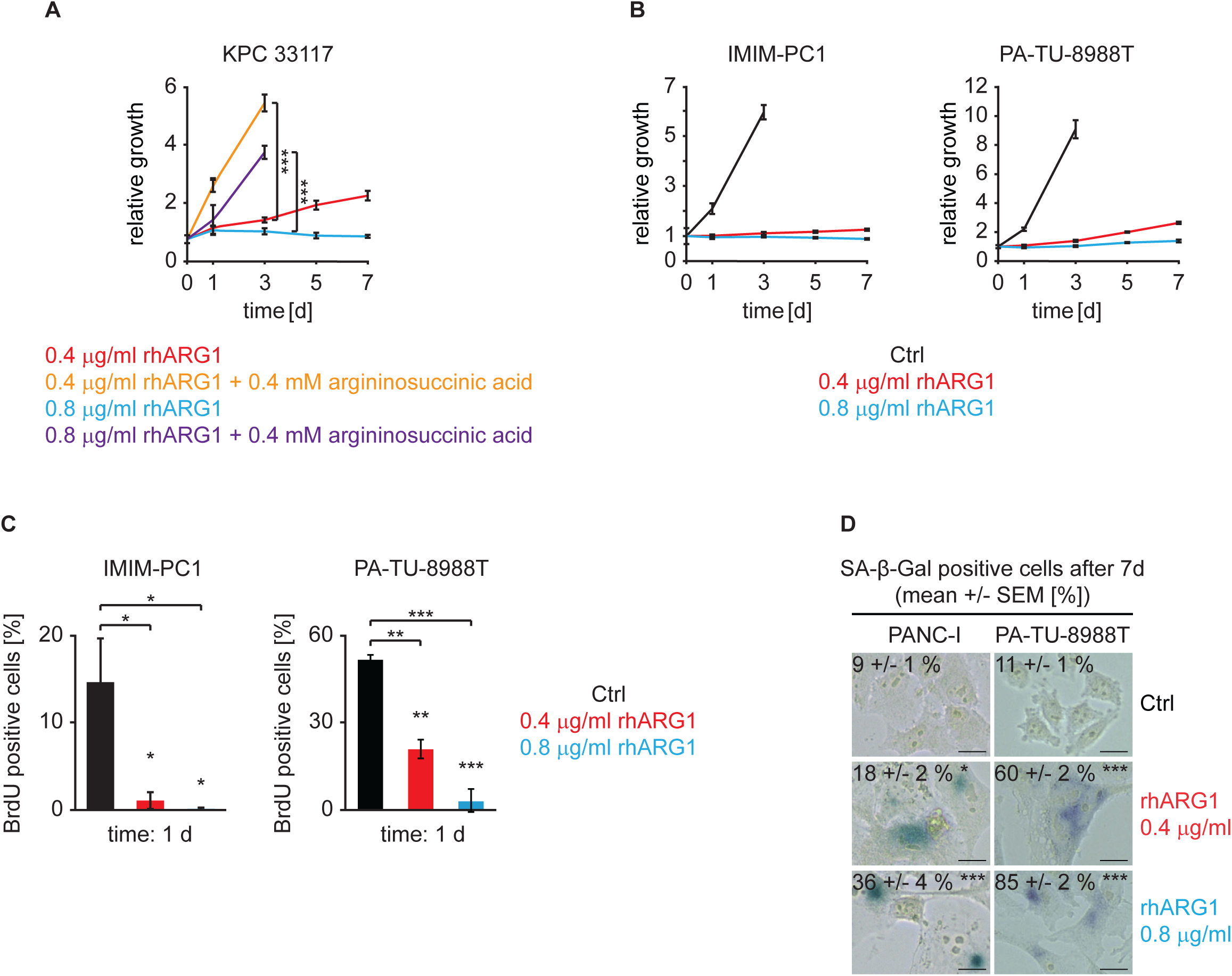
Murine arginine depleted PDAC cells resume proliferation in the presence of argininosuccinic acid and induction of senescence in human PDAC cells. A) Relative cell growth of murine PDAC cells treated as indicated. **B)** Relative cell growth of human PDAC cells upon PEG-rhARG1 treatment. **C)** BrdU-incorporation upon PEG-rhARG1 treatment in human PDAC cells. **D)** SA-β-Gal activity in human PDAC cells treated as indicated.

**Supplementary Figure 3:**
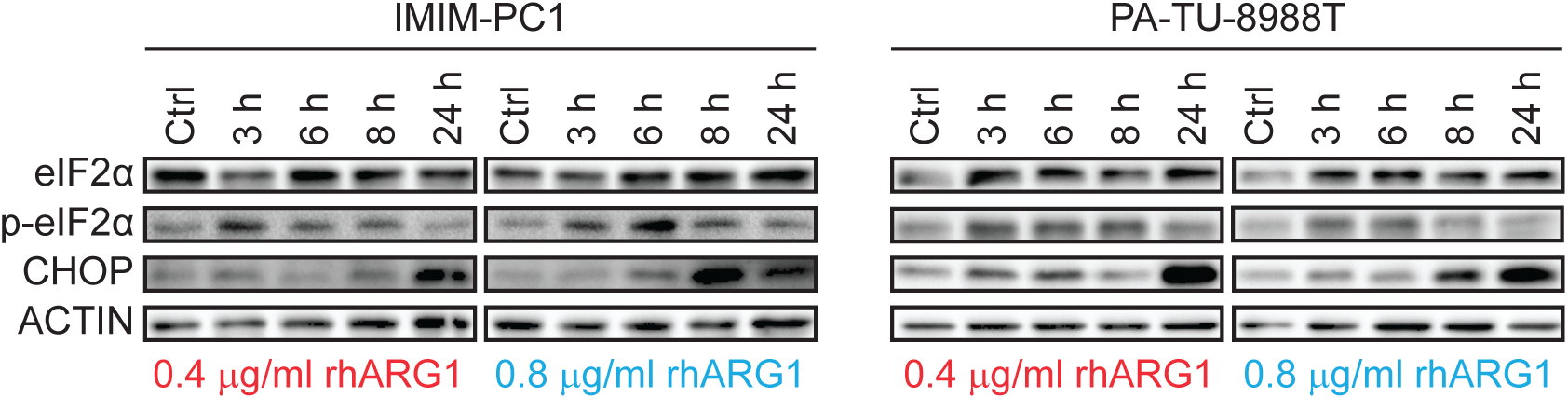
PEG-rhARG1 treatment activates the ISR in human PDAC. Immunoblotting of eIF2α, p-eIFα, CHOP and ACTIN as indicated.

**Supplementary Figure 4:**
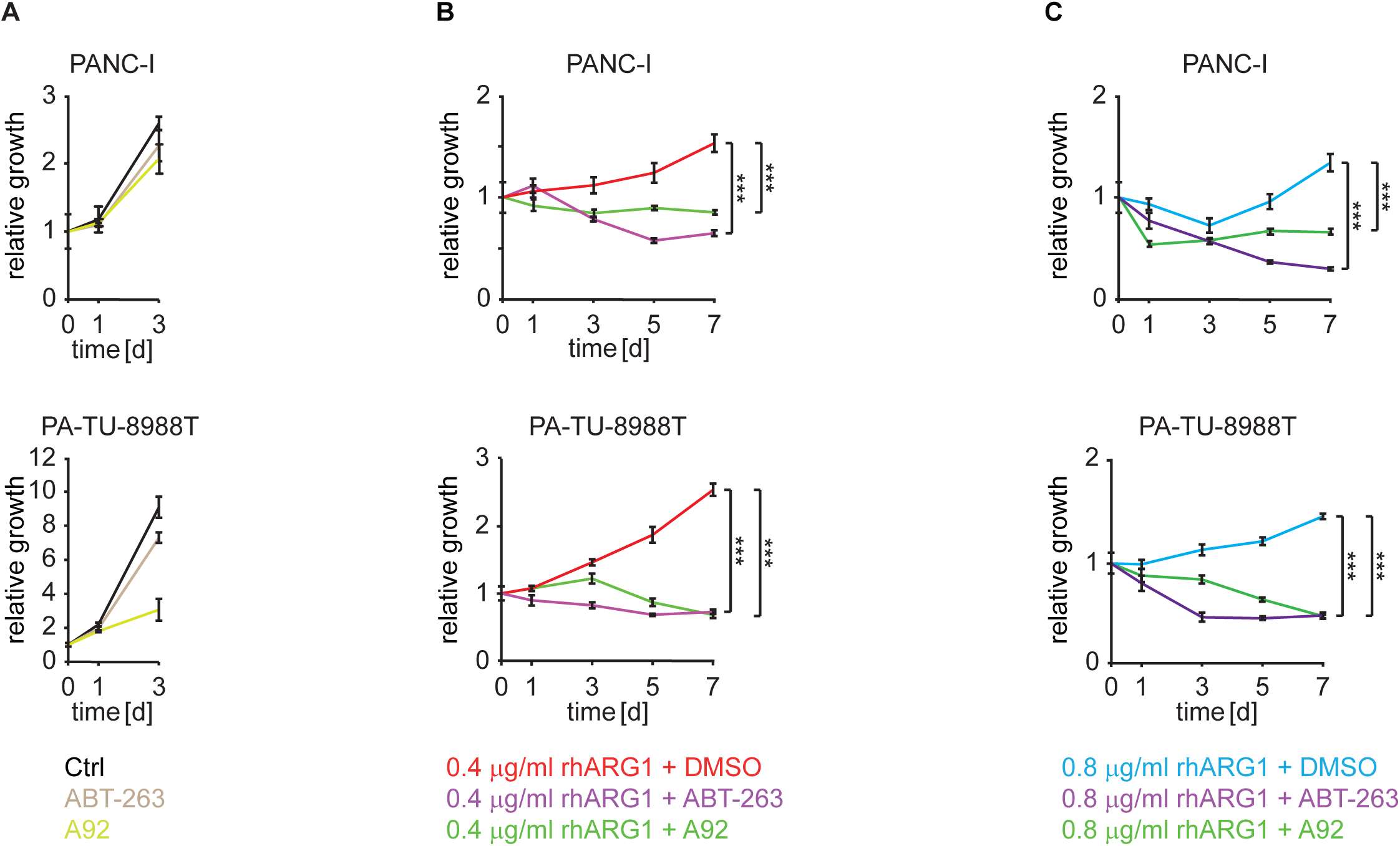
Synthetic lethality in human PDAC cells. A-C) Relative cell growth of human PDAC cells in response to the indicated treatments. Statistics: Student’s t test. ***P < 0.001.

## Supplementary Methods

### Chemical reagents

rhArgI-PEG5000 (BCT-100) was provided by Bio-Cancer Treatment International Limited, ABT-263/Navitoclax (#HY-10087) and A92/GCN-IN-1 (#HY-100877) were purchased from MedChemExpress.

### Cell Lines

Murine PDAC cell lines were generated by Jennifer Morton (Cancer Research UK Beatson Institute, United Kingdom) from *Pdx1-Cre; KRasG12D/wt; Trp53R172H/wt* (=KPC) mice. The number after KPC denotes the mouse number from which the cell was generated. Authenticated human cell lines were gifted by Dieter Saur (Technical University of Munich, Germany).

### Cell Culture

Cells were maintained under standard cell culture conditions (37 °C, 20% O2, 5% CO2) in regular cell culture medium (DMEM or RPMI supplemented with 10% FCS, 4.5 g/l glucose, 1 mM L-glutamine, 0.11 g/l pyruvate and 1% Penicillin-Streptomycin).

### Tissue Micro Array & Immunhistochemistry

The tissue micro array contained 101 individual human PDAC-specimen, was generated by Wilko Weichert (Technical University of Munich, Germany) and described previously [1]. The following antibodies were used to detect expression of ASS1 and OTC by immunohistochemistry following general protocols described in [2]: ASS1 mouse monoclonal (Thermo Fisher Scientific, MA5-17033), OTC rabbit polyclonal (Thermo Fisher Scientific, PA-5-35324)

### Immunoblotting

Immunoblotting was performed as previous described^1^. Briefly, cells were seeded at equal cell densities in regular cell culture medium 24 hours prior to treatment. At the end of the experiment, cells were lysed for 5 min with CytoBuster Extraction Reagent (Merck, 71009) supplemented with protease inhibitor (Thermo Scientific, A32953). The lysate was centrifuged in a tabletop centrifuge at maximum speed (16000 rpm) for 5 min at 4 °C. 10- 20 µg of protein were mixed with 5X SDS-loading buffer (250 mM Tris pH 6.8, 10% SDS, 50% Glycerol, 0.5 M DTT, 0.5 % Bromphenol blue), loaded onto a 15 % acrylamide gel and transferred onto PVDF membrane. The following antibodies were used ACTB/ACTIN BETA mouse monoclonal (Thermo Fisher Scientific, MA5-15739, 1:1000), ASS1 mouse monoclonal (Thermo Fisher Scientific, MA5-17033, 1:1000), ASL mouse monoclonal (Santa Cruz, sc-374353, 1:1000), ARG1 rabbit polyclonal (Thermo Fisher Scientific, PA5- 29645, 1:1000), CASP3/clCASP3 rabbit polyclonal (Cell Signaling, #9662, 1:500), CDK4 rabbit monoclonal (Absource, A5189, 1:1000), CHOP mouse monoclonal (Cell Signaling, #2895, 1:1000), CYCLIN D1 rabbit polyclonal (Cell Signaling, #2922,1:1000), eIF2α rabbit monoclonal (Cell Signaling, #5324, 1:1000), phospho-eIF2α rabbit monoclonal (Cell Signaling, #3398, 1:1000), GCN2 rabbit polyclonal (Cell Signaling, #3302, 1:500), OTC rabbit polyclonal (Thermo Fisher Scientific, PA-5-35324, 1:1000), P21 rabbit monoclonal (Bimake, A5163, 1:500).

### Immunofluorescence

Cells were seeded onto glass coverslips 24 hours prior to treatment. After fixation with ice-cold methanol/aceton (1:1) for 10 min at -20 °C, cells were washed and incubated in blocking solution (1% BSA in PBS) for one hour. Cells were then washed three times with PBS and incubated with primary antibody at 4 °C overnight. After three more washes with PBS, cells were incubated with secondary antibody for 1 hour at room temperature in the dark. DAPI was used as a nuclear stain. The following antibodies were used: BrdU mouse monoclonal (Cell Signaling, #5292, 1:1000), IncuCyte clCASP3/7 reagent (Essen Bioscience, #4440), H3K9me3 rabbit polyclonal (Abcam, ab8898, 1:500), HMGA2 rabbit polyclonal (Abcam, ab97276, 1:250), P16 rabbit polyclonal (Absource, #A5869, 1:50). Imaging of H3K9me3, HMGA2 and P16 was performed using an EVOS FL Auto 2 Imaging System (Fisher Scientific) and ImageJ software for image analysis.

### BrdU incorporation

Cells were seeded in 96-well plates 24 hours prior treatment. After treatment with 100 µM BrdU for 1 h, cells were fixed with icecold methanol for 5 min, followed by permeabilization with 0.2% Triton X-100 in PBS for 10 min at room temperature and incubation in 2 M HCl for 4 min. Cells were then incubated in blocking solution (3% BSA in PBS) for 30 min and then in primary antibody (Cell Signaling, #5292) diluted in 3% BSA in PBS overnight at 4 °C. After three washes with PBS, cells were incubated with secondary antibody for 1 hour at room temperature in the dark. Hoechst 33342 was used as a nuclear staining.

For measureing the BrdU incorporation, the Operetta® High-Content Imaging system (PerkinElmer) was used. 15 image fields per well were acquired with the 20x long WD objective and analysis was performed with the Harmony® High-Content Imaging and Analysis software (PerkinElmer). Nuclei were identified via Hoechst staining, nuclear BrdU intensity was measured and a threshold for BrdU positive nuclei was defined.

### Cell growth assay (SRB assay)

Cells were seeded in regular cell culture medium 24 hours prior treatment. Cells were fixed with 3.3% trichloroacetic acid solution for at least 1 hour at 4 °C. After washing cells three times with water and air-dried, the SRB staining solution (0.4% Sulforhodamine B sodium salt in 1% acetic acid solution) was applied onto the cells for 30 min. To remove unbound dye, cells were washed three times with 1% acidic acid. Stained cells were dissolved and diluted in 10 mM Tris pH 10.5 and absorbance at 510 nm is linearly correlated to the number of cells.

### RNA-seq (GSEA)

Cells were seeded in regular cell culture medium 24 hours prior treatment with 0.8 µg/ml rhARG1 for 3, 5 and 7 d. RNA was isolated using the RNeasy Mini Kit (Qiagen, #74104) in accordance with the manufacturer’s instructions. Equal amount of polyA selected RNA was used to generate sequencing libraries with the NEBNext® UltraTM Directional RNA Library Prep Kit for Illumina® (NEB #E7490) following the manufacturer’s instruction. Library size selection was performed using Agencourt® AMPure® XP Beads (Beckman Coulter, Inc.#A63881) and the resulting libraries were subjected to single end sequencing on NextSeq500 HighOutput flowcell for 75 cycles. Basecalling was performed using Illumina’s Basespace platform and the resulting FASTQs were aligned to the mm9 genome. Differential gene expression was performed using edgeR (Robinson et al., Bioinformatics, 2010) and the resulting table of counts was used to perform gene set enrichment analysis (Subramanian et al., PNAS, 2005).

### Senescence-associated β-galactosidase staining

At the end of the experiment, cells were washed with PBS (with 2mM MgCl_2_) and fixed for 15 min with freshly prepared fixation solution (4% paraformaldehye, 0.25% Glutaraldehyde). Cells were then repeatedly washed again (PBS (with 2mM MgCl_2_)) and incubated in staining solution (PBS with 2mM MgCl_2_, 1x KC solution (20x stock: 100 mM potassium ferricyanide (III), 100 mM Potassium hexocanoferrate (II)-Trihydrate in PBS), 1x X-Gal solution (40x stock: X-Gal in Dimethylformamide) for 15-24 hours at 37 °C in the dark without CO_2_. The pH of PBS/MgCl2 and the staining solution was adjusted to pH 5.5 for murine cells and pH 6.0 for human cells. After incubation, cells were washed with PBS three time and imaging was performed using EVOS FL Auto 2 Imaging System (Fisher Scientific) and ImageJ software for image analysis.

### Mass spectrometry

Metabolite extraction: Cells were washed with cold 154 mM ammonium acetate, snap frozen in liquid nitrogen and scraped off after addition of 0.5 mL ice-cold methanol/water (80/20, v/v) containing 0.25 μM of the internal standard lamivudine (Sigma). The resulting suspension was transferred to a reaction tube, mixed vigorously and centrifuged (2 minutes, 16 000g). Supernatants were transferred to a Strata® C18-E column (Phenomenex) which were previously activated with 1 ml of CH3CN and 1 ml of MeOH/H2O (80/20, v/v). The eluate was evaporated in a vacuum concentrator. The resulting residue was dissolved in 50 μl 5 mM NH4OAc in CH3CN/H2O (25/75, v/v). The pellets were kept for protein determination.

Liquid chromatography coupled mass spectrometry (LC-MS): Samples were diluted 1:2 with CH3CN and 5 μl of each sample was applied to a HILIC column (Acclaim Mixed-Mode HILIC-1, 3μm, 2.1*150 mm). Metabolites were separated at 30°C by LC using a DIONEX Ultimate 3000 UPLC system and the following solvents: Solvent A consisting of 5 mM NH4OAc in CH3CN/H2O (5/95, v/v) and solvent B consisting of 5 mM NH4OAc in CH3CN/H2O (95/5, v/v). The LC gradient program was: 100% solvent B for 1 minute, followed by a linear decrease to 40% solvent B within 5 minutes, then maintaining 40% B for 13 minutes, then returning to 100% B in 1 minute and 5 minutes 100% solvent B for column equilibration before each injection. The flow rate was maintained at 350 μL/min. The eluent was directed to the hESI source of the Q Exactive mass spectrometer (QE-MS) from 1.85 minutes to 18.0 minutes after sample injection (Thermo Scientific, Bremen, Germany). The scan range was set to 69.0 to 1000 m/z with a resolution of 70,000 and polarity switching (negative and positive ionization). Peaks corresponding to the calculated metabolites masses taken from an in-house metabolite library (MIM +/- H+ ± 2 mmU) were integrated using TraceFinder software (Thermo Scientific, Bremen, Germany).

Statistical analyses were performed using MetaboAnalyst 5.0 [3]. PCA 2D scoreplots, heatmaps (Distance measure: Euclidean, Algorithm: Ward) and dendrograms (Distance measure: Euclidean, Algorithm: Ward) were exported in svg format. Significantly changed metabolites were selected using ANOVA.

### RT-qPCR

Cells were seeded in regular cell culture medium 24 hours prior treatment. Total RNA was isolated using TRIzol (Invitrogen #15596018) in accordance with the manufacturer’s instructions. For RT-qPCR, 1 µg RNA was reverse-transcribed using the High-Capacity cDNA Reverse Transcription Kit (Thermo Fisher Scientific, #4368814). Using SYBR Select Master Mix for CFX (Thermo Fisher Scientific, #4472953), the reactions were run in triplicates in a total volume of 20 µl on the CFX96 Touch Real-Time PCR Detection system (BioRad). Each relative expression was normalized to 18S as a housekeeping gene. The following primers has been used 18S (fw: gtaacccgttgaaccccatt, rev: ccatccaatcggtactagcg), Il-1β (fw: acatcagcacctcacaagca, rev: tcttggccgaggactaagga), Il- 6 (fw: agccagagtccttcagagagat, rev: gagagcattggaaattggggt), Mmp-3 (fw: tccaggtgttgactcaaggg, rev: ttccaactgcgaagatccact).

### Statistics

All tests to define statistical significance were done with IBM SPSS statistics software and are stated in the figure legends. *P<0.05, **P<0.01, ***P<0.001

